# ELUCIDATION OF THE ROLE OF miRNA 4263 IN DYSREGULATED MITOCHONDRIAL ENERGETICS AND CARCINOGENESIS

**DOI:** 10.1101/2024.04.25.591199

**Authors:** Ashutosh Kumar Maurya, TV. Sruthi, V.B. Sameer Kumar

## Abstract

Dysfunctional mitochondria have been reported to be associated with several pathological conditions and in cancer, dysregulated mitochondrial metabolism is considered as an important hallmark of the disease. Cancer cells alter their mitochondrial machinery and activate glycolytic pathway as an alternate source of continuous energy, required for their indefinite growth. This modulation of the mitochondria could be due to the dysrupted expression of important mitochondrial genes involved in the normal functioning of the mitochondria. MicroRNAs are known to regulate the expression pattern of a variety of genes. With our in-silico analysis, we found that miR 4263 has targets on important mitochondrial genes, involved in mitochondrial energetics. Next, we checked the role of miR 4263 in modulating the mitochondrial metabolism and impact of this alteration on carcinogenesis. The results revealed that miR 4263 contributes to carcinogenesis in hepatic cells by altering the mitochondrial energetics.

## Introduction

MicroRNAs are small non-coding RNA molecule of ∼ 22 nucleotides (Shoubin Jhan et.al, 2020). MicroRNAs possess the ability of regulating the expression pattern of a variety of genes involved in normal functioning of the cells, thus plays a very important role in cells survival (Lyudmilla et.al, 2016). MicroRNAs are coded by nuclear DNA as well as mitochondrial DNA (Isabelle D. et.al, 2021).

Various reports suggests that nuclear coded microRNAs are localised in to the mitochondria and regulate the expression of target mitochondrial genes (Chiara Giordani et. al, 2021). Thus, nuclear coded microRNAs play important role in the normal functioning of the mitochondria, along with their mitochondrial counterpart, as they too possess the targets on mitochondrial protein coding genes (Goud et.al, 2015).

MicroRNAs, capable of changing the functioning of mitochondria, are called MitomiRs (Bandiera S., 2013). These MitomiRs either target internal proteins that are directly involved in ATP generation or mitochondrial membrane proteins that are involved in ATP transfer outside the mitochondria (Das S. et.al, 2012, Purohit P.K., et.al, 2021), so they play important role in various diseases associated with mitochondrial malfunctioning including cancer (Bienertova-Vasku J. et.al, 2013).

Dysregulated energetics and altered mitochondrial metabolism are significant cancer hallmarks (Sheng-Fan Wang et.al, 2023). In case of cancer, the mitochondrial machinery is altered and glycolytic pathway is activated to meet the energy requirement to support the infinite cellular growth (Narayanswami Badrinath et.al, 2018). Several studies have linked the altered mitochondrial metabolism with a variety of cancers (Fan S et.al, 2019).

Mitochondrial machinery is affected either by critical gene mutations or by silencing of genes involved in normal mitochondrial functioning (Evanthia Pangou et.al, 2021, Chaojun Y et.al,2019).

MicroRNAs could be a key player in the suppression of expression pattern of the crucial mitochondrial genes involved in electron transport chain (ETC), thereby altering the rate of ATP generation (Wee Lin Tan et.al, 2023). Several malignancies have been reported to have altered mitochondrial machinery together with elevated amounts of oncogenic microRNAs (Sheng-Fan Wang et.al, 2023). MicroRNA 4263 is an important oncomiR reported to be found at higher levels in exosomes derived from hypoxic tumor colony of HCC and plays an important role in carcinogenesis by inducing angiogenesis, an important hallmark of cancer (Sruthi TV, 2019).

This study deals with elucidation of the role of miR 4263 in carcinogenesis and modulation of mitochondrial functioning.

## Methodology

### Cell culture

Hela (cervical cancer cell line), WRL-68 (Human hepatic non cancerous cell line) and HepG2 cells (Hepatic carcinoma cell line) were cultured in DMEM supplemented with 10% FBS, antibiotic-antimycotic solution and L-Glutamine. The cells were maintained under standard culture conditions at 37°C with 5% CO2 and 95% humidity. For experiments, seeding density of 0.4 × 10^4^ cells (96 well), 0.6×10^6^ cells (30mm dish), 0.8×10^6^ cells (60mmdish), 2.2×10^6^ cells (100 mm dish) were used.

### miR 4263 cloning

miRNA 4263 was cloned in pCMV miR vector between BamH1 and Xho1 restriction sites and successful cloning was confirmed by sequencing.

### Transformation

The competent cells (DH5 *α*) were transformed with miR 4263 plasmid by heat shock method where the plasmid was incubated with the competent cells followed by a quick heat shock at 90°C for 2 minutes and then immediately transferring it on ice. The transformed cells were then plated on agar plate containing kanamycin.

### Plasmid isolation

A Single colony was picked from the agar plate and grown in the LB broth containing Kanamycin. The broth was incubated at 37°C in a shaking incubator. The plasmid was isolated by using Himedia midi kit following manufacturers protocol.

### Transfection

HeLa cells were seeded in 6 well plates and grown in a monolayer. After reaching 70% confluency, the cells were transfected with miR 4263 plasmid using PEI reagent and incubated for 24 hours in a CO_2_ incubator. After 6 hours of the incubation, the medium was replaced with fresh media and further incubated for 24 hours. Following this, the cells transfected with plasmids having fluorescent tags were observed under fluorescent microscope to check the efficiency of transfection

### Isolation of mitochondria

The mitochondria were isolated from the transfected cells using hypotonic buffer, where the cells were allowed to swell in the buffer for 10 minutes and then break open the cells to release the mitochondria. The cell suspension was then centrifuged at 1300g to remove the cell debris, followed by centrifugation at 12000g to get the mitochondrial pellet. The mitochondrial pellet was suspended in the mitochondrial resuspension buffer.

#### Sonication

The mitochondrial pellet was mixed with Lysis buffer and sonicated for 2 minutes at 70% amplitude with 15 sec ON and 30 sec OFF cycle on 4°C. The solution obtained, was centrifuged at 12000g for 10 minutes. The supernatant was collected and protein estimation was done followed by sample preparation for SDS PAGE.

#### Protein estimation

Protein level of mitochondria was estimated by Bradford method (Bradford etal,1976). To achieve this, 10μl of sample and 90μl of bradford reagent (50 mg Coomasie Brilliant Blue-G250 in 25ml ethanol and 50ml of phosphoric acid made upto 100ml with water) was added in triplicates in 96 well plate and the absorbance was taken at 595nm by multimode plate reader. The concentration of protein was calculated from the standard plot to BSA with concentration range from 10μg-100μg.

#### SDS-PAGE

Protein sample was prepared by mixing of 6x SDS loading dye and boiling it at 90°C for 10 minutes in water bath. The sample was immediately kept on ice and briefly centrifuged before loading on SDS-PAGE gel. The electrophoresis was carried out by using Bio-Rad electrophoresis unit. The protein samples were run through the stacking gel at 80V for 15 minutes and through the resolving gel at 100V at room temperature until the dye reached the end of the gel.

### Western blot analysis

The purity of the mitochondrial pellet was checked by western blot using mitochondria specific antibody (VDAC). Also, the mitochondrial pellet was checked for the nuclear and cytoplasmic contaminants using Histone H3 antibody for Nucleus and Hexokinase HK3 antibody for the cytoplasm.

### RNA Isolation

RNA was isolated from the mitochondrial pellet as well as from the total cell using trizole reagent. Following this, the concentration of the RNA was checked by using nano drop.

### Polyadenylation of RNA

Poly A tail was added to the RNA by Poly A Polymerase enzyme, using manufacturers protocol. This reaction set up was incubated at 37°C for 30 minutes followed by heat inactivation for 5 minutes at 65°C.

### cDNA synthesis

The polyadenylated RNA were used for the synthesis of miRNA 4263 specific cDNA by Kang method. Apart from this, total RNA was used to synthesize the cDNA for checking the expression of mitochondrial genes.

### Real Time PCR (qRT PCR)

Quantative real time PCR was performed to check the expression pattern of the microRNA 4263 and other mitochondrial genes in mitochondria before and after over expression of miR 4263.

### mRNA stability assay

To elucidate the targeting of mitochondrial genes by miR 4263, mRNA stability assay was performed. The cells were transfected with miR 4263 using PEI method. 24 hours post transfection the cells were treated with actinomycine D at 0, 1, 3, 6 and 12 hours. The samples were collected at each time point for gene expression study.

### Oxygraph analysis

To check the phenotypic effects of the down regulation of the mitochondrial genes by miR 4263, the oxygraph analysis was performed, where the oxygen consumption level was checked in the miR 4263 over expressed samples and compared with the control samples. In brief, the cells were grown in a 6 well plate and transfected with the candidate microRNAs using PEI method. After 48 hours of incubation at 37°C, the cells were trypsinized and the cell pellet was resuspended in respiration buffer. Later, 1ml of the cellular suspension was added to oxymeter and oxygen intake reading was recorded for 10 minutes. The readings were used to plot the graph to represent the oxygen consumption by the mitochondria.

### Exosome isolation

The exosomes were isolated from the miR 4263 over expressing cells using PEG method. To achieve this the cells were transfected with miR 4263 and media was replaced with serum free medium. 48 hours post transfection; the spent media was collected and centrifuged at 2000g for 1/2 hour, to remove cell debris. Following this, the cells were mixed with PEG solution in 1:2 ratios and incubated at 4 degree C overnight. Finally, the solution was centrifuged at 12000g for 1 hour. The exosomal pellet was dissolved in PBS and protein estimation was done.

### Cell migration Assay

Cancer cells have the metastatic properties, where they move from its origin to another place and form a secondary tumor. To mimic this in-vitro we perform the cell migration assay with an objective to check, if the exosomes from miR 4263 over expressing cancer cells could induce cellular migration of normal Hepatic cells. A scratch was made in the WRL monolayer and treated with exosomes isolated from miR 4263 over expressing cancer cells. The cells were allowed to fill the gap formed by the scratch for 48 hours and the images were taken 0, 24, 48 hours respectively and quantified using ImageJ.

### Soft Agar Colony formation assay

To elucidate the role of miR 4263 in inducing carcinogenic properties in normal heaptic cells, the colony formation assay was performed where the WRL cells were suspended in the low melting agarose and treated with the exosomes. The cells were incubated at 37°C for 28 days and allowed to form the colonies. The colonies were stained with Coomasie brilliant blue and images were taken from 20 different locations and quantification was done by using ImageJ.

### In-Silico analysis

miR 4263 target prediction analysis and scoring for mitochondrial genes was done using five algorithms i.e. MirBase, miRanda, Target Scan, miRDB and MicroRNA.org. Five highest scored mitochondrial genes (ND6, ATP6, Cyto-B, Cox1, ND4L) were selected for target validation.

#### Statistical analysis

All the data in the study were expressed as the mean with the standard error mean of at least three experiments, each done in triplicates. SPSS 11.0 software was used for analysis of statistical significance of difference by Duncan’s One way Analysis of Variance (ANOVA). A value of P<0.05 was considered significant.

## Results

### Cellular and mitochondrial levels of miR 4263 in cancerous and non-cancerous hepatic cell lines

As first part of the study, we checked the levels of microRNA 4263 in HepG2 and non-cancerous hepatic cell line (WRL) by qRT-PCR. The results revealed that, there was a higher level of miR 4263 in HepG2 cells when compared with the control cells. Following this, we checked the levels of miR 4263 in the mitochondria of cancerous & non-cancerous hepatic cells (i.e. HepG2 & WRL) and the results suggested that level of miR 4263 was found significantly lower in the mitochondria of the HepG2 cells when compared with the control cells (WRL), even though the total cellular levels miR 4263 was higher.

**(Figure 1)**

**Figure 1:**
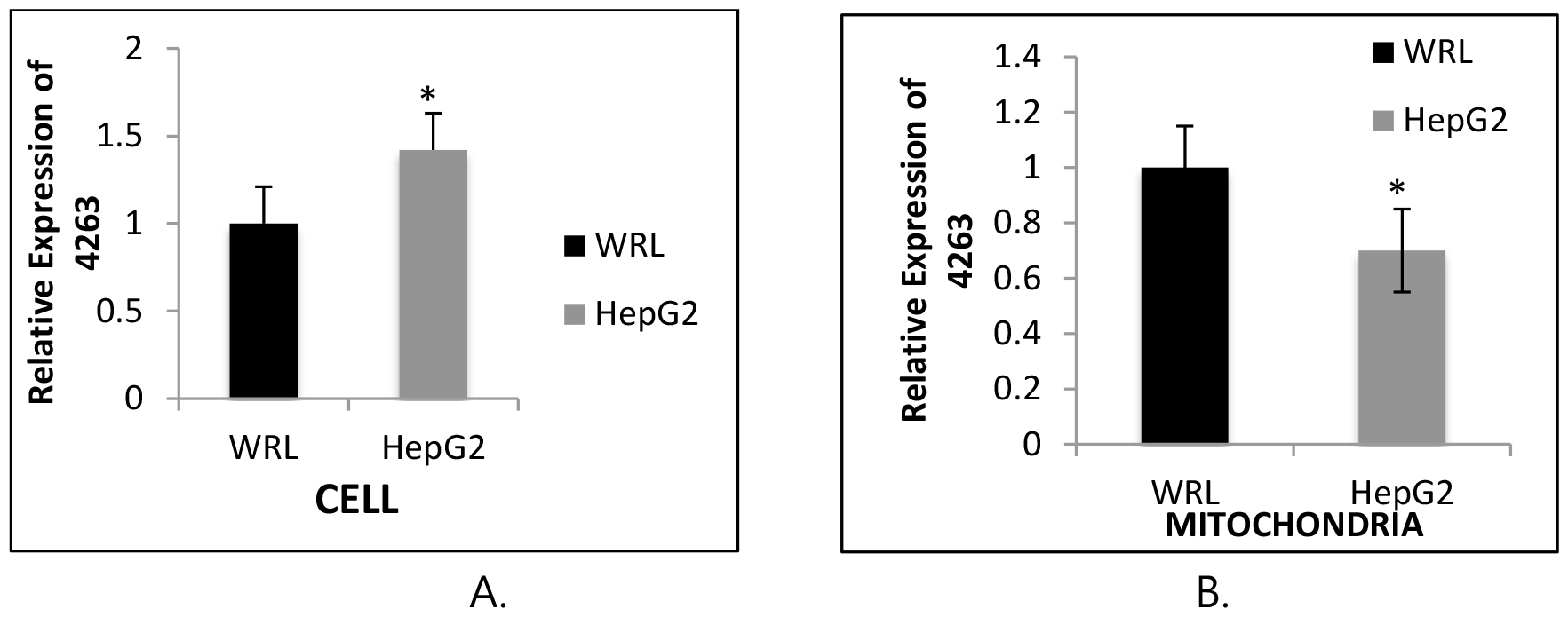
Differential expression pattern of miR 4263 in cytoplasm and mitochondria. RT PCR analysis was performed to check the levels of miR 4263 in mitochondrial and cellular fraction of HepG2 cells, keeping WRL as control. The results suggested that level of miR 4263 was significantly low in the mitochondria of HepG2 cells. A.) Relative expression levels of miR 4263 in the cells B.) Relative levels of miR 4263 in the mitochondria. Results presented are average of three experiments ± SEM each done at least in triplicate, p<0.05. *Statistically significant when compared to control.

### Over expression of miR 4263 resulted in its preferential targeting to the mitochondria

As we found that the microRNA 4263 was found significantly low in the mitochondria, we over expressed miR 4263 in HepG2 cells and checked its relative levels in the cytoplasm & mitochondria. The results revealed that, miR 4263 gets targeted to the mitochondria in a selective manner when compared with the mock transfected cells, with a 4.5 fold increase in mitochondria and 3 fold increase in the cytoplasm, when compared to the levels in the mock transfected cells.

**(Figure 2)**

**Figure 2:**
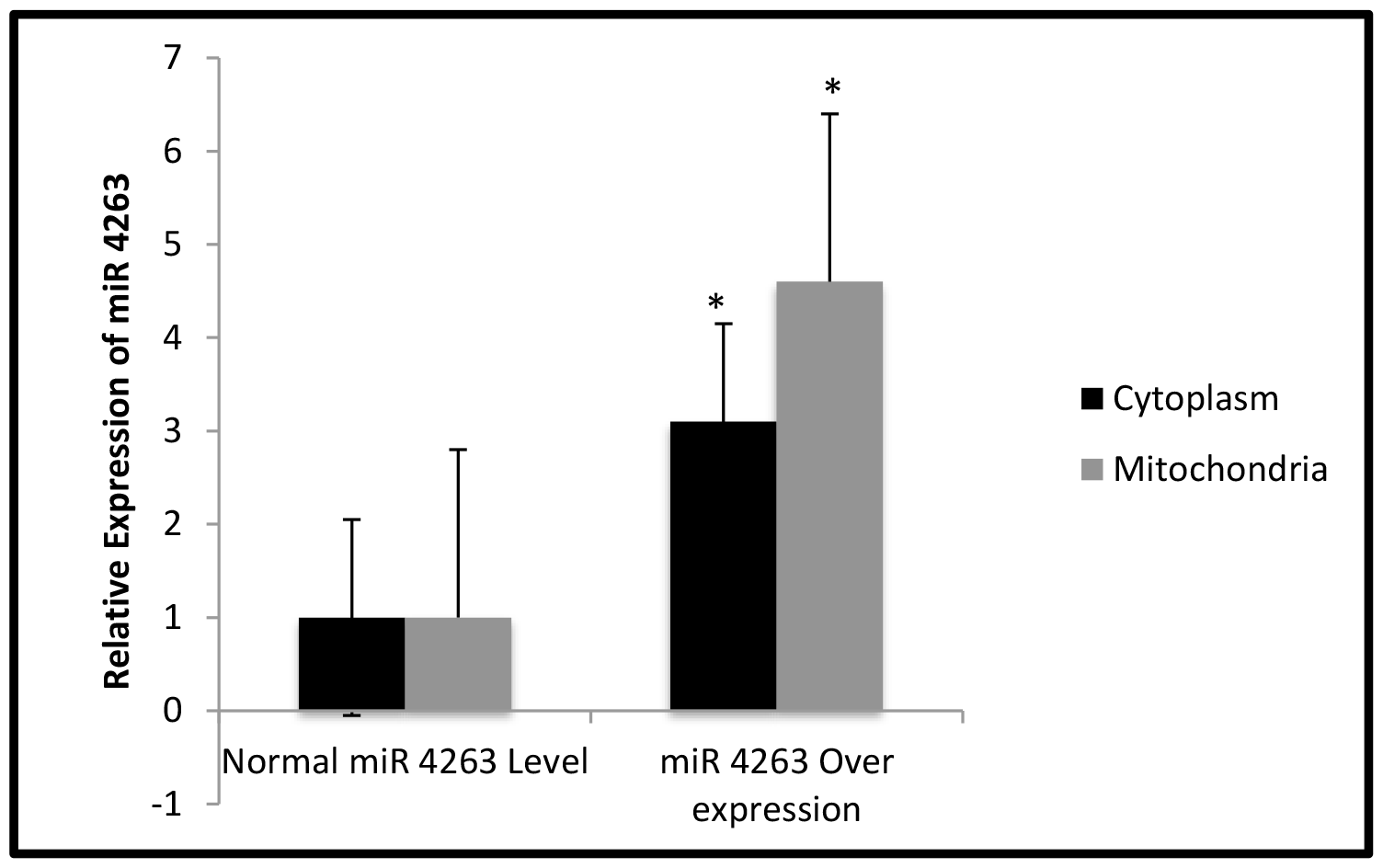
miR 4263 gets targeted to the mitochondria in a selective manner. miR 4263 was over expressed in HepG2 cells followed by purification of mitochondria & isolation of RNA from mitochondrial as well as cytoplasmic fractions and, qRT PCR was then performed to check the levels of miR 4263. The results revealed that miR 4263 gets targeted to mitochondria in a selective manner. Results presented are average of three experiments ± SEM each done at least in triplicate, p< 0.05.*Statistically significant when compared to control.

### miRNA 4263 target the mitochondrial genes

After confirmation of the localization of miR 4263 to the mitochondria, next we checked the effect of this enrichment on the expression level of mitochondrial target genes. To check this, we performed target validation study. The results of this study revealed that level of all the target genes of miR 4263 were significantly lowered when compared with the control cells, confirming our in-silico target prediction analysis.

**(Figure 3)**

**Figure 3:**
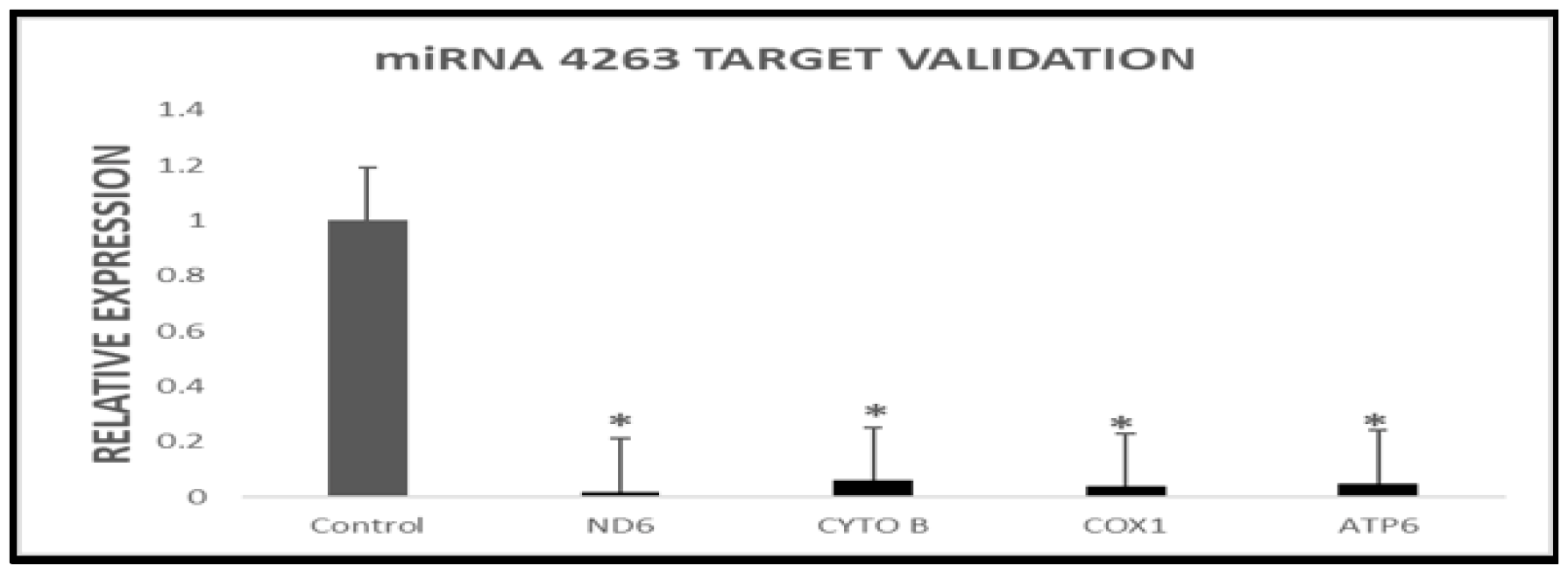
miR 4263 down regulates all its target genes. miRNA 4263 was over expressed in HepG2 cells and the mitochondria isolated. RT-PCR analysis was performed to check the impact of increased miR 4263 level on the expression pattern of target mitochondrial genes. The result revealed that the expression level of all the mitochondrial target genes of miR 4263 went significantly down. Results presented are average of three experiments ± SEM each done at least in triplicate, p<0.05.*Statistically significant when compared to control.

### mRNA stability assay revealed the targeting of ND6 gene by miR 4263

Target validation results revealed that levels of all the mitochondrial target genes were significantly lowered by miR 4263. To further check the targeting of mitochondrial genes by miR 4263, we performed mRNA stability assay and the results revealed that level of all the target genes went down, upon miR 4263 over expression, most effectively ND6, suggesting that 3’ UTR of these genes harbour putative miR 4263 binding sites.

**(Figure 4)**

**Figure 4:**
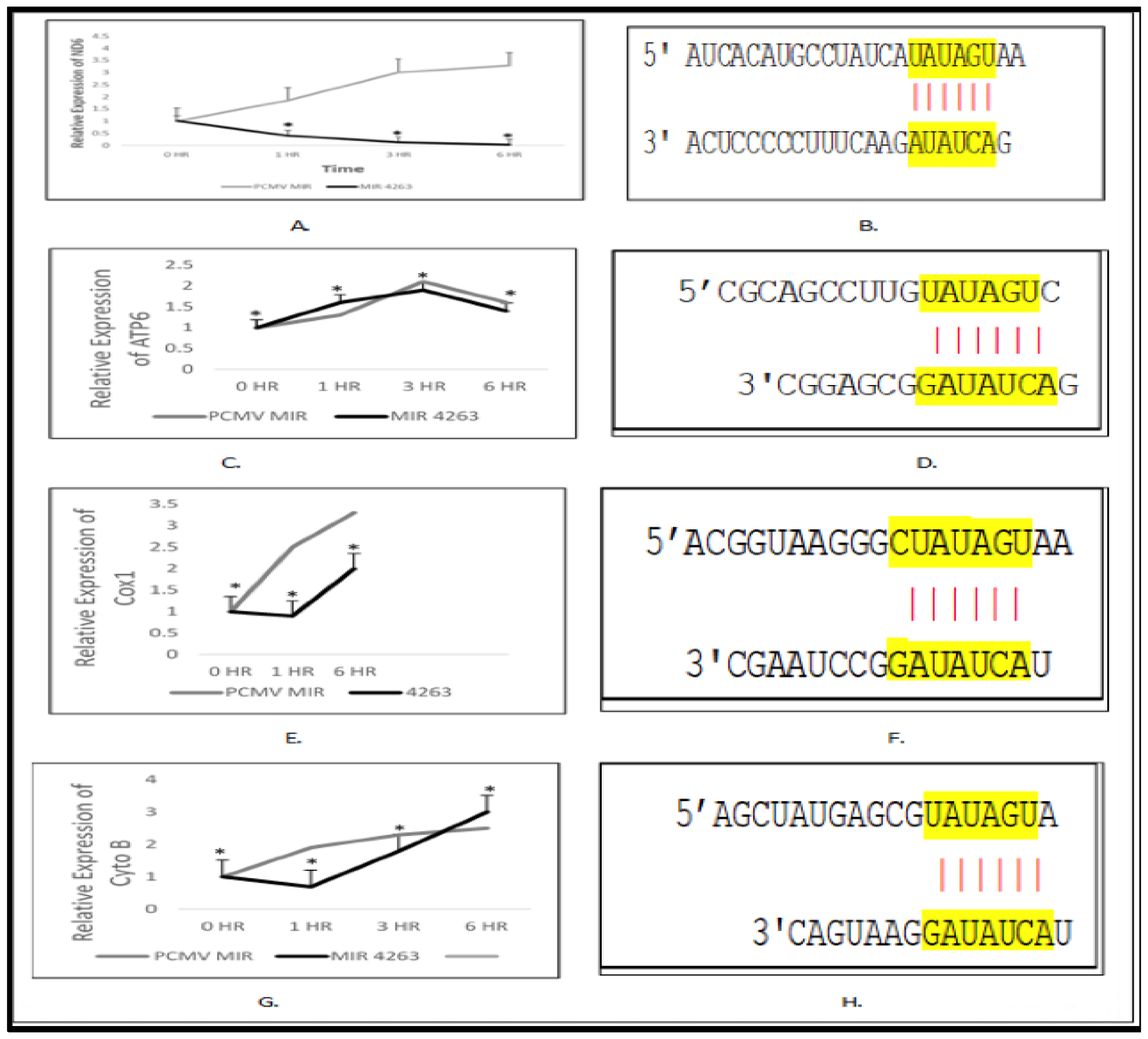
3’ UTR analysis and mRNA stability assay revealed the direct targeting of ND6 by miR 4263. miR 4263 was overexpressed in HepG2 cells and 24 hours post transfection, the cells were treated with actinomycin D and RNA samples were collected at 0, 1, 6 and 12 hours respectively. Following this, RT-PCR analysis was performed to check the expression pattern of target genes. Results revealed that expression of ND6 went significantly down with time suggesting its direct targeting by miR4263. The expression pattern of Cox1, ATP6 and Cyto-B also was altered, but not significant. A.) Relative expression of ND6 B.) 3’ UTR analysis of ND6 C.) Relative expression of ATP6 D.) 3’ UTR analysis of ATP6 E.) Relative expression of Cox1 F.) 3’ UTR analysis of Cox1 G.) Relative expression of Cyto-B H.) 3’ UTR analysis of Cyto-B. Results presented are average of three experiments ± SEM each done at least in triplicate, P<0.05.*Statistically significant when compared to control.

### Oxygen consumption by mitochondria of HepG2 cells went significantly lower in miR 4263 over expressing cells

The mRNA stability assay suggested that miR 4263 targets the ND6 gene which is very crucial for complex 1 of electron transport chain in the mitochondria. Any alteration in the expression level of this gene may dysregulate the normal functioning of the mitochondria. So, to analyze the phenotypic effect of the down regulation of this gene on the mitochondrial metabolism, we performed the oxygraph analysis, where we checked the levels of the oxygen consumption by the HepG2 cells.

The result of the oxygraph analysis revealed that the levels of oxygen consumption by the HepG2 cells went significantly down upon miR 4263 over expression when compared with the mock transfected controls, suggesting its role in modulating the mitochondrial machinery.

**(Figure 5)**

**Figure 5:**
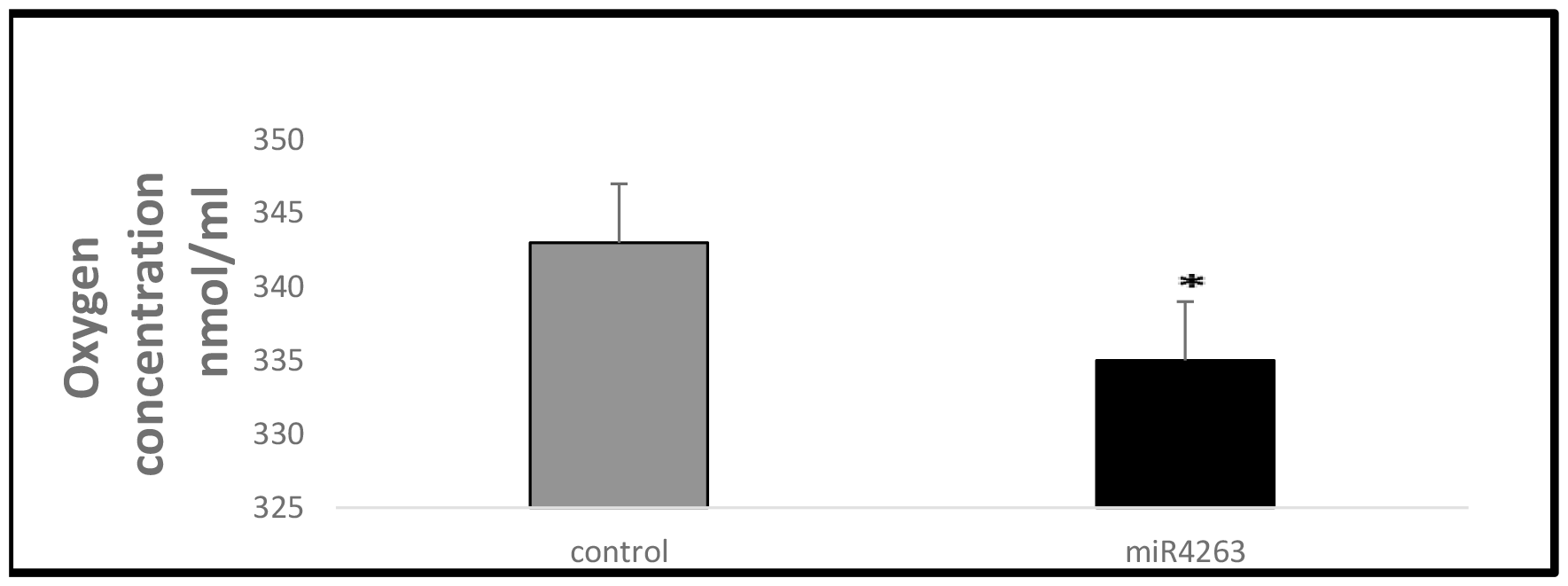
Levels of oxygen consumption by the cells went significantly down, when miR 4263 was over expressed. miR 4263 was over expressed in HepG2 cell line and 24 hour post transfection, cells were harvested and oxygraph analysis was performed. The results revealed a decrease in O_2_ consumption by the mitochondria of miR 4263 overexpressing cells when compared with the mock transfected cell. Results presented are average of three experiments ± SEM each done at least in triplicate, p< 0.05.*Statistically significant when compared to control.

### MicroRNA 4263 increases the migratory properties of cancerous and non-cancerous hepatic cells

From our target validation and mRNA stability assay results, we found that miR 4263 gets targeted to the mitochondria and down regulate the expression of ND6 gene important in complex 1 of ETC and further it reduced the oxygen consumption by the HepG2 cells, probably due to the altered mitochondrial metabolism. So, we next checked the impact of this altered mitochondrial metabolism in carcinogenesis. To achieve this, we performed the cell migration assay using non cancerous & cancerous hepatic cells (WRL & HepG2) and treated it with the exosomes isolated from miR 4263 over expressing cancer cells or mock transfected Hela cells.

The results of cell migration assay revealed an increase in the rate of cellular migration when treated with exosomes isolated from the miRNA 4263 over expressing cells, when compared to the cells treated with the exosomes isolated from the mock transfected cells and the pattern was found to be persistent in both, cancerous as well as non-cancerous hepatic cell lines.

**(Figure 6 & Figure 7)**

**Figure 6:**
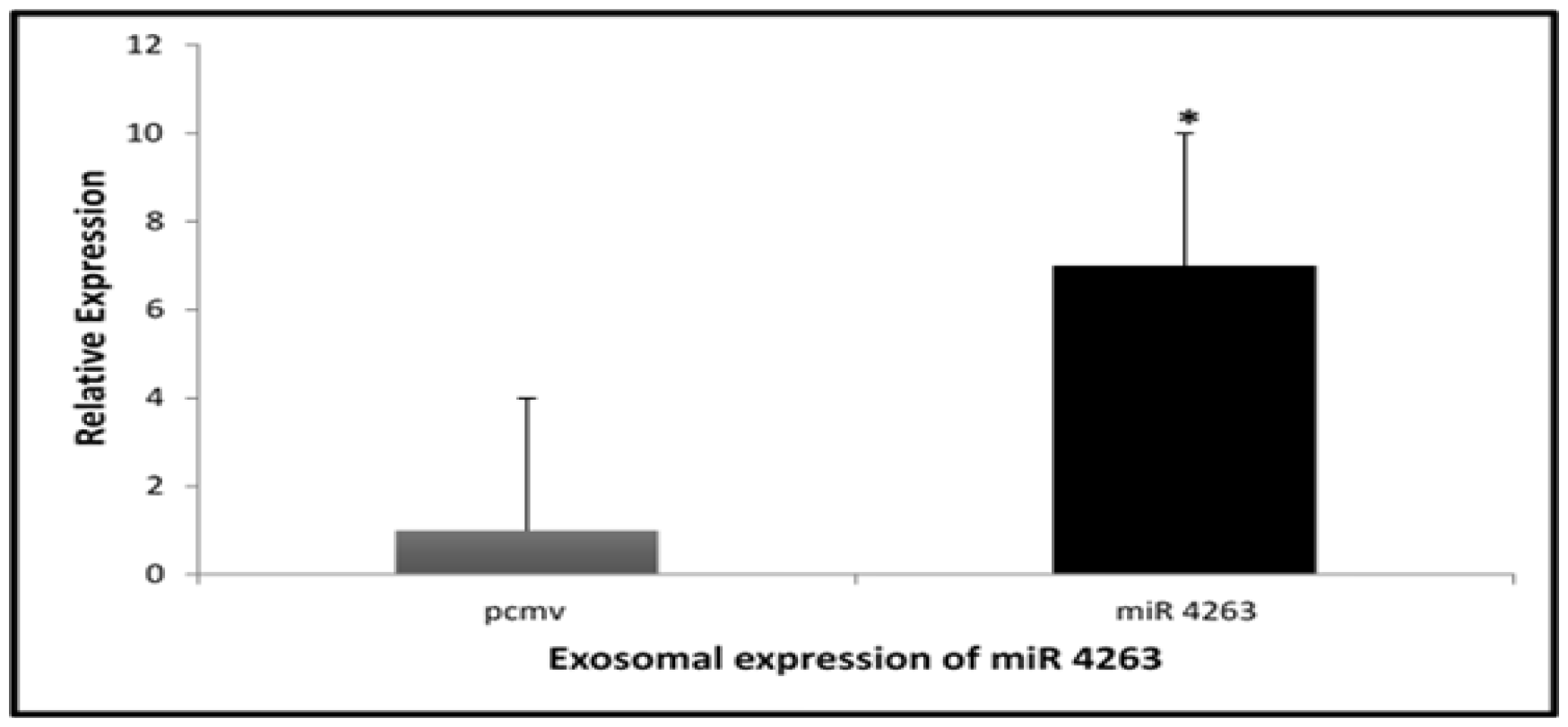
Exosomes isolated from miR 4263 over expressing cells were enriched with miR 4263. Real time PCR analysis was performed to check the enrichment of miR 4263 in the exosomes isolated from miR 4263 over expressing cells and compared with the exosomes isolated from the mock transfected control cells. The results revealed that exosomes isolated from microRNA 4263 over expressing cells, shown 7 fold higher level of microRNA 4263. Results presented are average of three experiments ± SEM each done at least in triplicate, p<0.05.*Statistically significant when compared to control.

**Figure 7:**
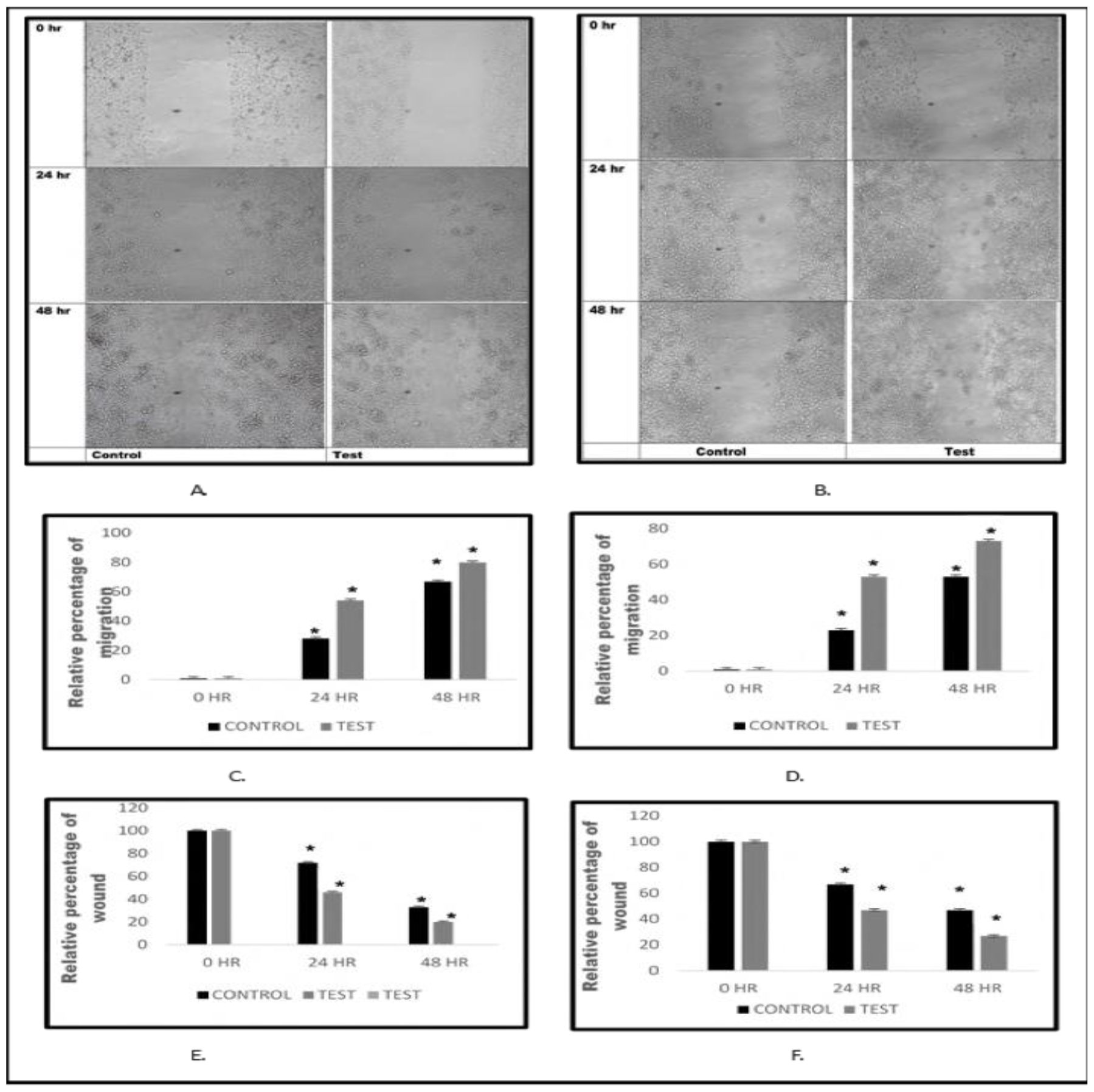
miR 4263 escalates the cellular migration when treated with exosomes isolated from microRNA 4263 over expressing cells. Cells were grown in a monolayer and a scratch was made, followed by exosome treatment. The microphotographs were taken at 0, 24, and 48 hours. The distance/gap covered by the cells with time was estimated by image-J software. The result revealed that the rate of the cellular migration got escalated when treated with exosomes enriched with miR 4263. A.) Microphotograph of cell migration pattern (HepG2) with respect to time B.) Microphotograph of cell migration pattern (HepG2) with respect to time C.) Relative percentage of migration by HepG2 cells, at 0, 24 & 48 hours D.) Relative percentage of migration by WRL cells at 0, 24 & 48 hours E.) Relative percentage of wound healing at 0, 24 & 48 hours (HepG2). F.) Relative percentage of wound healing at 0, 24 & 48 hours (WRL). Results presented are average of three experiments ± SEM each done at least in triplicate, p<0.05.*Statistically significant when compared to control.

### Exosomes from miR 4263 over expressing cells enhances colonization property

We performed colony formation assay to check the colonogenic ability of miR 4263 in normal and cancerous hepatic cells. Here, we treated hepatic cell colonies with the exosomes isolated from miR 4263 over expressing cells or mock transfected control cells. Colony formation assay result showed an increase in the number and size of the colonies when treated with the exosomes enriched with miR4263 as compared with the colonies treated with exosomes isolated from the mock transfected cells.

**(Figure 8)**

**Figure 8:**
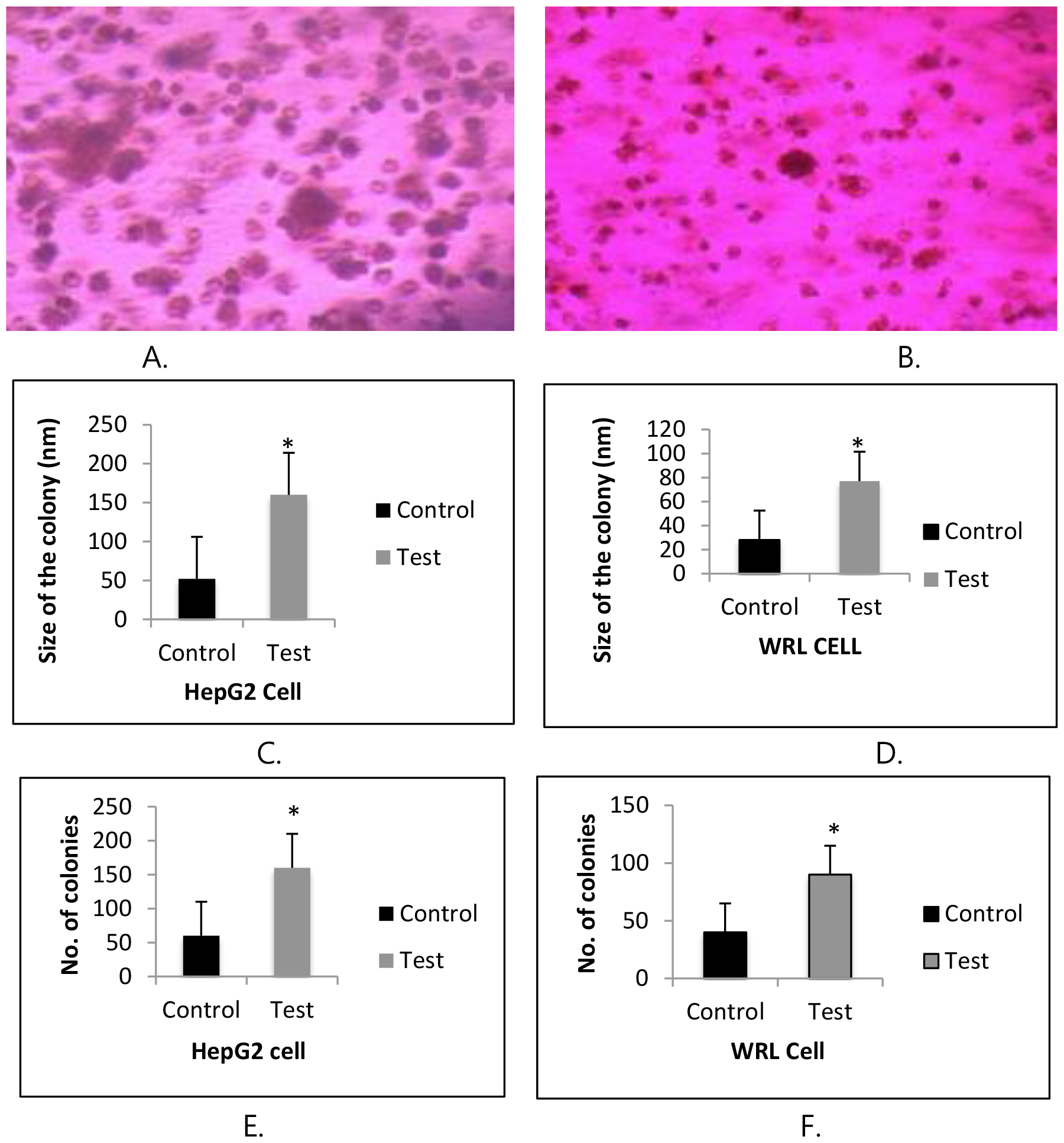
miR 4263 enhances the tumor size when treated with exosomes isolated from microRNA 4263 over expressing cells. HepG2 and WRL colonies were treated with exosomes isolated from miR 4263 overexpressing cells and allowed to grow for 28 days. Following this, microphotographs were taken at 20 different regions and 20 colonies from each region was taken for the size estimation by image-J software. The number of colonies were counted manually from 20 regions to estimate the number of tumor colonies. The results revealed that the size as well as the number of colonies got enhanced in HepG2 and WRL cells, when treated with exosomes enriched with miR4263. A.) Representative microphotograph of HepG2 cell colony B.) Representative microphotograph of WRL cell colony C.) Comparative colony size of HepG2 cells. D.) Comparative colony size of WRL cells E.) Number of colonies of HepG2 cells, when compared with control F.) Number of colonies of WRL cells, when compared with control. Results presented are average of three experiments ± SEM each done at leastin triplicate, p<0.05. *Statistically significant when compared to control.

## Discussion

Dysregulated mitochondrial energetics has been established as one of the major hallmark of the cancer (Shan-Millan I, et.al, 2023). By modifying mitochondrial metabolism, cancer cells change how they produce energy in more advanced stages of the disease. (Sheng-Fan Wang et.al, 2023). In a variety of cancer, the glycolytic pathway has been reported as an alternate mode of energy, employed by the cancer cells for their exponential growth. (Ganpathi K et.al, 2013).

The modulation of mitochondrial machinery could either result due to the mutations in important genes involved in mitochondrial metabolism or due to the suppression of expression of the genes engaged in the energy production (Amanda Lopes, 2020). The suppression of mitochondrial activity could be brought in by the action of the microRNAs (Zhang 2021). The microRNAs are small non-coding RNAs, capable of regulating the expression pattern of a variety of the genes involved in various cellular functions (Kioomars S. et. al, 2019). MicroRNAs which could alter the mitochondrial metabolism are called mitomiRs (Isabelle D. et.al, 2021). MitomiRs can either be coded by mitochondrial genome or nuclear genome. From the cytoplasm, nuclear coded miRNAs are transported to the mitochondria where they function by controlling the expression levels of metabolism related mitochondrial genes, resulting in reprogramming of the mitochondrial machinery and replacement of electron transport chain with glycolytic pathway (Purohit P.K., et.al, 2021).

Hepatocellular carcinoma (HCC) is 3rd most common type of cancer, with more than 1 million estimated yearly cases by the 2025 (Josep M et.al, 2021). The key role of various microRNAs including mitomiRs, have been reported to be important in the initiation and progression of HCC (Yi Fu et.al, 2019, Zhang Lisheng et.al, 2013). MicroRNA 4263 has been found at higher levels in the exosomes isolated from hypoxic tumor colonies of HCC, and was found to be involved in the process of angiogenesis, (Sruthi TV, 2019), thereby acting as a key player in carcinogenesis.

The main objective of this study was to check the role of miR 4263 in the modulation of the mitochondrial metabolism and its impact on carcinogenesis. Since miR 4263 is a nuclear coded miRNA, first we checked its level in the mitochondria of HepG2 by qRT-PCR. The results revealed that miR 4263 was significantly low in the mitochondria. As established by the various studies, that nuclear coded miRNAs gets localized to the mitochondria, next we over expressed miR 4263 in HepG2 cells and qRT-PCR results suggested that miR 4263 got enriched in the mitochondria by 4.5 folds in a preferential manner.

Our bioinformatic results revealed that miR 4263 has targets on important mitochondrial genes, and also we found that miR 4263 gets targeted to the mitochondria, so next we performed target validation study to check if miR 4263 could alter the expression of mitochondrial genes upon reaching the mitochondria. The qRT-PCR results revealed that, miR 4263 significantly lowered the expression levels of all mitochondrial target genes. Next, we performed mRNA stability assay for all mitochondrial target genes, to further check their targeting by miR 4263 and the results revealed that ND6 is most effectively targeted by miR 4263, whereas Cox1, CytoB and ATP6 gets targeted however to non significant extent and the result falls in line with the 3’ UTR analysis that revealed that 3’ UTR of ND6 harbour putative miR4263 binding sites. Since, ND6 gene plays important role in electron transport chain and is crucial for the mitochondrial machinery, its down regulation could directly impact the mitochondrial metabolism.

So, next we checked the phenotypic effects of this down regulation by performing the oxygraph analysis, to check the oxygen consumption by mitochondria of miR 4263 overexpressed cells and found that oxygen consumption level went significantly down, suggesting its role in altering the functioning of mitochondrial machinery by targeting ND6, which is involved in complex 1 of electron transport chain. Several studies have established the role microRNAs in improving the migratory properties of hepatic cancer cells, e.g. miR 221 has been found to promote HCC cells migration by targeting Plant homeo domain finger 2 (PHF2), a tumor suppressor, regulating P53 (Yi Fu et.al, 2019) and miR-665 promotes proliferation and migration of HCC cells by targeting PTPRB involved in hippo signalling (Yuanchang Hu et.al, 2018). MicroRNAs have also been found in escalating the colonogenic properties of HCC, for example microRNA 657 has been reported to promote tumorigenesis in HCC by targeting transducing-like enhancer protein-1 (TLE-1) (Zhang Lisheng et.al, 2013), and also, miR-125b has been suggested to be involved in HCC progression (Dorothy Fan et.al, 2012).

Since, these studies demonstrated that under over expressed conditions, the microRNAs promoted cell migration and colony formation, so next we checked if the over expression of miR 4263 too, can induce carcinogenesis in hepatic cell lines. To achieve this, we performed cell migration assay and colony formation assay by treating the hepatic cells with the exosomes isolated from miR 4263 over expressing cells. The results suggested that migratory and colony formation ability of hepatic cells got enhanced when treated with the exosomes enriched with miR 4263.

These results therefore suggest that miR 4263 possess carcinogenic ability and that it possibly involve the modulation of mitochondrial metabolism.

## Acknowledgment

We acknowledge Indian Council of Medical Research, Ministry of Health, Govt. of India for the financial assistance in form SRF and Kerala state council for Science Technology & Environment, Govt. of Kerala for fellowship in the form of JRF and SRF to Mr. Ashutosh K. Maurya. We also acknowledge Central University of Kerala for providing all the necessary facilities to carry out this research work.

## Author Contributions

The authors confirm contribution to the paper as follows: Study conception and design: VBSK, Bioinformatics and wet lab work: AKM. Cloning of miR 106b: STV. All authors reviewed the results and approved the final version of the manuscript.

## Conflicts of Interest

The authors declare that they have no conflicts of interest to report regarding the present study.

## SUPPLEMENTARY DATA

## Notes

### Competing Interest Statement

The authors have declared no competing interest.

